# ModPhred: an integrative toolkit for the analysis and storage of nanopore sequencing DNA and RNA modification data

**DOI:** 10.1101/2021.03.26.437220

**Authors:** Leszek P. Pryszcz, Eva Maria Novoa

## Abstract

**Motivation:** DNA and RNA modifications can now be identified using Nanopore sequencing. However, we currently lack a flexible software to efficiently encode, store, analyze and visualize DNA and RNA modification data.

**Results:** Here we present *ModPhred*, a versatile toolkit that facilitates DNA and RNA modification analysis from nanopore sequencing reads in a user-friendly manner. *ModPhred* integrates probabilistic DNA and RNA modification information within the FASTQ and BAM file formats, can be used to encode multiple types of modifications simultaneously, and its output can be easily coupled to genomic track viewers, facilitating the visualization and analysis of DNA and RNA modification information in individual reads in a simple and computationally efficient manner.

**Availability and Implementation:** *ModPhred* is available at https://github.com/novoalab/modPhred, is implemented in Python3, and is released under an MIT license.

**Supplementary Data:** Supplementary Data are available at *Bioinformatics* online.

## 1. INTRODUCTION

Third generation sequencing technologies have revolutionized our ability to identify base modifications in single molecules (Novoa *et al.*, 2017; Kelleher *et al.*, 2018; Garalde *et al.*, 2018; Q. Liu, Fang, *et al.*, 2019; Loman *et al.*, 2015). While many tools have been developed in the recent years to detect DNA and RNA modifications from nanopore sequencing datasets (Yuen *et al.*, 2020; Stoiber *et al.*, 2017; Ni *et al.*, 2019; Leger *et al.*, 2019; H. Liu *et al.*, 2019; Jenjaroenpun *et al.*, 2021; Q. Liu, Georgieva, *et al.*, 2019; Pratanwanich *et al.*, 2020; Begik *et al.*, 2021), there are limited tools allowing retrieval, storage, manipulation and visualisation of modification information (De Coster *et al.*, 2020; Leger, 2020).

Currently, the only available algorithm to extract and store DNA or RNA modification information from basecalled FAST5 datasets is *megalodon* (https://github.com/nanoporetech/megalodon), a tool developed by Oxford Nanopore Technologies (ONT) that relies on a previously trained basecalling model to extract methylation information from each raw Fast5 read, which is then dumped into a plain text file that will contain all predicted modified sites. However, *megalodon* presents several caveats and limitations: (i) it only supports m^5^C and m^6^A DNA modification detection, (ii) it cannot be used with direct RNA sequencing datasets that are mapped to the genome, (iii) it does not integrate modification information within the FastQ format, (iv) it does not have the ability to encode multiple RNA modification types simultaneously (e.g. m^5^C and hm^5^C), (v) it cannot be parallelized by splitting the input FAST5 files into separate read chunks, and (vi) it does not offer options for downstream analyses or visualization of the results (**Table S1**).

Here, we present *ModPhred*, a toolkit that encodes DNA and/or RNA modification information within the FastQ and BAM formats, allowing its analysis and visualization at single molecule resolution (**Fig. 1A**). We show that *ModPhred* can extract and encode modification information from basecalled FAST5 datasets 4-8 times faster than *megalodon*, while producing output files that are 50 times smaller (**Table S2**). Finally, we illustrate the applicability of the *ModPhred* toolkit for the analysis of both DNA and RNA modifications. The toolkit is easy to use by the non-bioinformatic expert, and generates user-friendly reports to facilitate the downstream analyses as well as several forms of visualization of the modification information (**Fig. 1B**), both at per-site as well as at per-read level.

**Figure 1.**
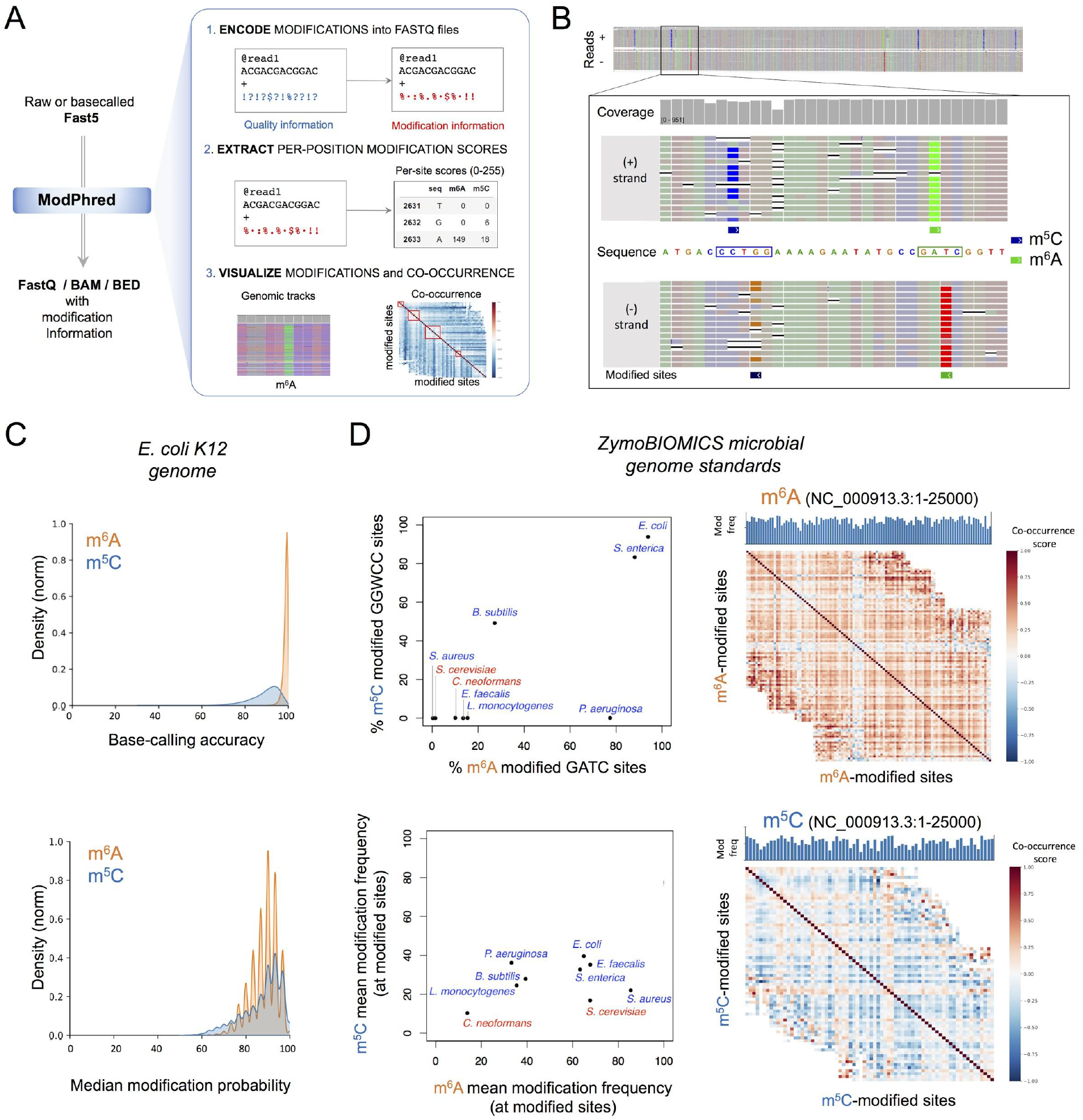
Overview of *ModPhred*. **(A)** Schematic representation of ModPhred input, output and steps performed. Briefly, *ModPhred* uses as input raw or basecalled Fast5, and returns FASTQ, BAM and BEDGraph with modification information. To achieve this, *ModPhred* first encodes modification information into FASTQ files (modEncode) substituting the quality information, and then into the BAM files (modAlign). *ModPhred* can then easily extract modification information from BAM files to generate reports (modReport). Finally, *modPhred* can be used to visualize the results (modAnalysis). See also Figure S1 for additional details on each of the 4 individual modules of *ModPhred*. **(B)** IGV visualization of BAM files generated using *ModPhred*. Since *ModPhred* stores modification information in the base quality field, per-read modification information can be visualized in IGV browser by coloring reads based on per-base quality information. **(C)** Density plots of basecalling accuracies (upper panel) and median modification probabilities (lower panel) at predicted modified sites, generated by *modReport*. See also Figure S2. **(D)** Analysis of ZymoBIOMICS microbial reference using *ModPhred*. In the left panels, global analysis of m^6^A and m^5^C modification levels across different species are shown. In the right panels, cooccurrence analysis of m^6^A (upper panel) and m^5^C (lower panel) DNA modifications are depicted, for the same genomic region (NC_000913.3_1-25000).

## 2. MATERIALS AND METHODS

*ModPhred* is conceived to efficiently encode, process and visualize DNA and RNA modification data from nanopore sequencing datasets. *ModPhred* only requires as input a reference genome and reads in FAST5 format, which can be raw (non basecalled) or basecalled using RNA or DNA modification-aware basecaller (guppy version 3.4+). *ModPhred* extracts and integrates DNA/RNA modification information into the FASTQ and BAM files, by including: (i) information regarding the type of DNA/RNA modification (e.g. m^5^C or hm^5^C), and (ii) probability scores of the most probable modification, for each nucleotide in each read. By decoupling the processes of basecalling and modification annotation, *ModPhred* can rapidly extract the list of modifications without the need of recomputing the basecalling step.

*ModPhred* is subdivided into 4 modules, and performs the following tasks: (i) encoding of modification probabilities in FastQ, with optional basecalling step *(modEncode);* (ii) alignment of reads that includes generation of BAM files with modification information (*modAlign*); (iii) extraction of modification information from mapped reads *(modReport);* (iv) downstream analyses (*modAnalysis*), which include plotting of DNA/RNA modifications within genomic track viewers (*modPlot*), computing correlations between modified positions (*modCorrelation*) and read clustering based on modification patterns (*modCluster*) (**Fig. S1**, see also ***Supp. Methods***).

Firstly, *ModEncode* processes the FAST5 reads and stores the most likely type of modification for each base in every read. This is achieved by encoding the modification probability in the form of an ASCII character, replacing the basecalling qualities that are by default encoded in FastQ files (**Fig. S1**, see also ***Supp. Note 1***). Such storage of information results in data compression, simplicity and versatility. Since probability of modification is stored inside the FastQ file, no external databases or additional files are needed for calculation or visualisation of modifications. Per-base modification probabilities are also stored in BAM files that are derived from FastQ during the alignment step, which is performed by *modAlign. modReport* then calculates a list of statistics for every base of the genome and reports positions that are modified. Finally, *modAnalysis* generates graphical representations of modification statistics, as well as high-level analysis of DNA/RNA modification distributions, including co-occurrence of modifications and per-read clustering based on similarity of DNA/RNA modification patterns.

## 3. IMPLEMENTATION OF ModPhred

We first tested *ModPhred* for the annotation and analysis of DNA modifications in microbial datasets (**Table S3**). We should note that current *guppy* basecalling models (versions 3.2.1 and later) can so far only detect m^5^C in CCWGG and CpG contexts, and m^6^A in GATC contexts. Therefore, our analysis was limited to these modifications and sequence contexts. To this end, we analysed a high-coverage (900x) *E.coli* K12 DNA genome sequencing dataset. *ModPhred* reported 38,897 m^6^A and 29,165 m^5^C-modified positions in *E. coli* chromosome, with mean modification frequencies at the modified sites of 63.2% and 42.4% (**Fig. S2A-B**) and mean modification probabilities of 0.878 and 0.865 for m^6^A and m^5^C, respectively (**Fig. 1C**, see also **Table S4**). ModPhred reported 99.7% of *E. coli* GATC and CCWGG sites as ‘modified’ (i.e. modification frequency was greater than 0.05), whereas only 0.12% of CpG sites were reported as modified. The latter are expected to be false positives, since CpG methylation is not known to exist in *E. coli*. Similar results were observed in a second *E. coli* dataset with lower coverage (250x), showing high reproducibility across datasets (**Fig. S2A-B**, see also **Table S4**).

We then applied *ModPhred* to the ZymoBIOMICS microbial DNA reference dataset (**Table S3**). We find that m^6^A and m^5^C predictions, both in terms of modification frequency as well as in terms of penetrance, largely vary across species. *E. coli* showed the highest penetrance of m^6^A and m5C modifications in GATC and CCWGG sequence contexts, in agreement with previous results (**Fig. 1D**). Closely related species, such as *S. enterica*, showed similar penetrance of m6A and m5C modifications in GATC and CCWGG sequence contexts. However, the vast majority of species analyzed did not show high penetrance of m^5^C modifications in CCWGG sites, suggesting that either the penetrance in these species is either low, or that the motif in which m^5^C is embedded is different than CCWGG (**Fig. 1D**, see also **Table S4**).

Finally, we applied *modPhred* to direct RNA nanopore sequencing datasets. However, we should note that currently, there are no publicly available *guppy* models for the detection of RNA modifications. Thus, to illustrate the applicability of *modPhred* in direct RNA sequencing data, we employed in-house *taiyaki-trained* RNA modification-aware models that were trained using synthetic RNA molecules (see ***Supp. Methods***). Specifically, we examined the ability of *modPhred* to predict and annotate RNA modifications in different mixes of RNA-modified datasets, finding that *modPhred* accurately recapitulates the expected RNA modification frequencies (**Table S5**, see also **Fig S2C**). Moreover, we illustrate how modPhred can be used for per-read cluster analysis based on their RNA-modification profiles, illustrating its applicability to identify read populations with similar co-occurrence of RNA modification patterns (**Fig. S2D**). Overall, our results show that modPhred can be applied both for the analysis of DNA and RNA modifications in genomic and transcriptomic datasets.

### Benchmarking of modPhred and comparison to available tools

*ModPhred* runtimes were compared to *megalodon* on two publicly available genome sequencing datasets: (i) the *E.coli* DNA genome sequencing (PRJEB22772) and (ii) the ZymoBIOMICS microbial reference DNA genome sequencing (PRJNA477598) datasets (**Table S2**, see also **Fig. S3**). We observed that *ModPhred* was 4-8x faster than *megalodon*, while producing 50x smaller result files than *megalodon*. Moreover, we found that *megalodon* was poorly applicable to high coverage samples (PRJEB22772, *E. coli* sample with 900x coverage) as the process would not finish after 120 hours, limiting *megalodon’s* applicability in high coverage and/or large genomes (**Table S2**). By contrast, we observed that the runtime of *modPhred* scaled well both with coverage and genome size, and that it was mainly limited by the speed of basecalling process (**Tables S6-S8**). This limitation can be easily overcome by using a multi-GPU system as well as by processing each project or sample in smaller batches. Finally, we should note that *modPhred* can perform remote basecalling, allowing many remote clients to process read batches in parallel from multiple workstations (or computing cluster nodes). By contrast, *megalodon* is designed to process all reads from a given sample at once and in a single workstation equipped with one or more dedicated GPUs, which leads to decreased parallelization and increased computing times (**Table S3**, see also ***Supp. Note 2***).

## ACKNOWLEDGEMENTS

LPP is supported by funding from the European Union’s H2020 research and innovation programme under Marie Sklodowska-Curie grant agreement No. 754422. This work was partly supported by the Spanish Ministry of Economy, Industry and Competitiveness (MEIC) (PGC2018-098152-A-100 to EMN). We acknowledge the support of the MEIC to the EMBL partnership, Centro de Excelencia Severo Ochoa and CERCA Programme / Generalitat de Catalunya. We would like to thank the CRG Scientific Information Technologies (SIT) facility for setting up the GPU cluster and continuously upgrading the corresponding software that was needed to benchmark *modPhred* and *megalodon* using diverse guppy versions.

