## Supplementary Figure for "ModPhred: an integrative toolkit for the analysis and storage of nanopore sequencing DNA and RNA modification data"



**CONTENTS**


1. **SUPPLEMENTARY METHODS**
2. **SUPPLEMENTARY NOTES**
3. **SUPPLEMENTARY TABLES**
4. **SUPPLEMENTARY FIGURES**
5. **SUPPLEMENTARY REFERENCES**

### 1. SUPPLEMENTARY METHODS

#### Nanopore sequencing datasets

Nanopore DNA sequencing raw Fast5 were obtained from publicly available datasets (*E. coli*: PRJEB22772; Zymo mock community: PRJNA477598) from previously published studies (McIntyre *et al.*, 2019; Jain *et al.*, 2017). Direct RNA sequencing datasets with and without RNA modifications were obtained from publicly available datasets ($m^6A$: PRJNA511582; $m^5C$: PRJNA563591; $hm^5C$: PRJNA548268; UNM:PRJNA511582) (Liu *et al.*, 2019; Begik *et al.*, 2021).

#### Training RNA modification basecalling algorithms using *Taiyaki*

$m^5C$, $m^6A$ and $hm^5C$ RNA modification-aware basecalling models were obtained by training *taiyaki* (https://github.com/nanoporetech/taiyaki) using $m^5C$-modified, m6A-modified and $hm^5C$-modified *in vitro* transcribed sequences ('curlcakes'), as well as their unmodified counterparts. Briefly, *taiyaki* trains neural networks using nanopore current intensity signal information, and outputs models that are compatible with *guppy* basecaller. We followed the default *taiyaki* training workflow described in the GitHub repository. The trained model is available at https://github.com/novoalab/modPhred/tree/main/data. We should note that this trained model is not applicable to *de novo* basecall *in vivo* sequences, for two reasons: (i) 'curlcake' sequences were *in vitro* transcribed by replacing one of the canonical bases with modified base across all positions (ie $m^6A$ instead of A), and consequently, there are no instances of given modified base in the context of unmodified counterpart (e.g. AAAAA will be only represented as $m^6Am^6Am^6Am^6Am^6A$); and (ii) 'curlcakes' were designed to cover all possible 5-mers, but most likely they do not cover sufficient sequence diversity to train a deep learning basecalling model applicable to other sequences such as *in vivo* transcripts. As a result, our *taiyaki* model is not applicable to basecall and *de novo* predict RNA modifications transcriptome-wide.

#### FAST5 basecalling

DNA whole genome sequencing datasets were basecalled *on-the-fly* with *guppy* version 3.6.1 or 4.0.15. We should note that *ModPhred* v1.0 can be run on FAST5 that have been basecalled with *guppy* versions 3.4+, whereas *megalodon* v2+ can only be run with versions 4.0+. Both *modPhred* and *megalodon* can be run with Fast5 files that have been basecalled before or using *on-the-fly* basecalling. Direct RNA sequencing datasets were basecalled with modification-aware pre-trained *taiyaki* models.

Performing basecalling *on-the-fly* using *ModPhred* is the recommended option, since the basecalling process produces large output files that need to be parsed during modification encoding. This results in high I/O operation overhead, especially in the case of network filesystems and it is in general much slower than running basecalling *on-the-fly*. For example, basecalling PRJEB22772 with `--fast5_out` parameter took over 19 hours, while running the entire modPhred pipeline using on-the-fly basecalling took just over 5 hours (**Table S7**).

**Encoding modification probabilities in FastQ**

*ModPhred* can extract the list of modifications without the need of recomputing the basecalling step, thanks to encoding the per-read and per-nucleotide modification probabilities into the FastQ files, replacing the basecalling qualities. A major reason for encoding modification probabilities in the FastQ files is that accessing information from Fast5 files is slow compared to accessing information in FastQ files.

FastQ basecalled qualities are typically stored as 93 ASCII characters valued from 33 (`!`) to 126 (`~`). *Guppy* basecaller reports the probability of a base being modified in the Fast5 files, in the form of 8-bit integers scaled from 0 (no modification) to 255 (modification). These values are reported separately for all modifications that a given model has been trained for (**Figure S1A**), and must be converted into ASCII characters prior to embedding them into the FastQ files.

The encoding of modification information into ASCII characters will vary depending on the number of modifications that the user wants to store. The number modifications for which we store information can be adjusted by setting `--MaxModsPerBase` parameter. Theoretically, we are able to store up to 368 modifications (92 per base) directly in FastQ format given we limit the information about modifications status to binary form, meaning either base is carrying one of the 368 modifications or not (**Figure S1A**). By default, *modPhred* stores information from 3 modification types for every base (A, C, G, T/U), making up to 12 modifications in total. In this default scenario ( `--MaxModsPerBase 3`), the modification probability ASCII scores encoded into the FastQ basecalled qualities will be computed as follows:

$$Q = 255 * \max\{p_1, p_2, p_3\} / 31 + 31 * M_i$$

where $M_i$ is: 0 if first, 1 if second or 2 if third modification of given base has the highest probability of modification (p).

All metadata, including modification names, symbols and number modified of bases, are stored in the Fast5 directory as `mod_data.pkl`.

**Detection of genome/transcriptome positions with modifications**

Alignments in *ModPhred* are performed using *minimap2*, and sorted BAM files are stored in `out_dir/minimap2` directory. For direct RNA samples, splice-aware mapping is automatically performed by *modPhred*, which means that it can process reads mapped both to the genome as well as to transcriptome. Following the alignment of the reads, *modPhred* calculates the following statistics for every reference position: (i) stranded depth of coverage at a given position (number of aligned reads reported for +/- stand separately); (ii) basecalling accuracy (frequency of most common base ignoring indels), (iii) modification frequency (frequency of every modification), and (iv) median modification probability (median probability across modified bases).

By default, *modPhred* reports above metrics into a gzipped file (`mod.gz`) only for sites with modifications fulfilling the below criteria (relevant parameters are given in brackets):

- alignments with mapping quality below 15 are ignored (`--mapq 15`),
- at least 25 reads aligned at given positions (`--minDepth 25`),
- at least 5% of reads being modified for this position (`--minModFreq 0.05`)
- bases with modification probability above 50% are considered as truly modified (`--minModProb 0.5`).

**Quality filtering, visualisation and analysis of DNA and RNA modification results**

*ModReport* generates violin (`mod.gz.svg`) and pairwise/density (in `plots/` directory) plots for depth of coverage, basecall accuracy, modification frequency and median modification probability of bases with modifications for all detected modified positions from all samples (**Fig. 1C**, see also **Fig. S1** and **S2B,C**). In addition, *modReport* generates bedGraph files with modified positions for every sample, which can be visualised using a genome browser (**Fig. 1B**). The modification status can be directly visualised in genome browsers if the user chooses to color the reads based on their per-base qualities, since modification status is encoded as basecall quality. Finally, *ModAnalysis* can be used to facilitate the analysis of predicted DNA/RNA modifications, by generating 3 different types of plots: (i) venn diagrams, to examine the reproducibility of predicted DNA/RNA modifications (modPlot), (ii) co-occurrence heatmaps, to examine the co-occurrence of predicted DNA modifications within reads covering a given genomic regions (modCorrelation); and (iii) per-read clustering, to identify subsets of RNA reads with similar RNA modification patterns (modCluster).

**Code availability**

Source code and workflows to reproduce this work are available via *modPhred* GitHub repository https://github.com/novoalab/modPhred.

### 2. SUPPLEMENTARY NOTES

**Supplementary Note 1: Guppy basecaller API troubleshooting**

While working on this project, we have encountered numerous difficulties related to Guppy basecaller API, known as *pyguppyclient* (https://github.com/nanoporetech/pyguppyclient).

Firstly, decreased efficiency in parsing the data was observed, largely due to the sequential processing of individual reads by the API. To process one read, the API submits it to the basecalling server, waits until the read is fully basecalled and retrieves the results from the server. We found this process to be very slow (~4 reads per second), because the basecalling server is idle during read submission and result retrieval, while the API process is idle during read basecalling. This can be sped-up using multiple API connections, but we observed saturation when using 8-12 connections (~40 reads per second), most likely due to multiple queries to the server from each API instance. To overcome this limitation, we opted to submit reads in batches (for example 100 reads each) and gradually retrieve reads that were already basecalled. Such implementation greatly (~100x) speeds up API (~400 reads per second) because the basecalling server is fully utilised (since there are no idle times between individual read submissions). However, we should note that the API doesn't provide such functionality (batch submission), and we have implemented this functionality inside the script *guppy_encode_live.py*, which is part of *modEncode*. We should note that the user still has to install a *pyguppyclient* compatible with the guppy version of choice, as guppy_encode_live.py internally uses *pyguppyclient*. Detailed instructions on how to use and install the *pyguppyclient* to run *modPhred* can be found in the documentation (https://modphred.readthedocs.io).

Secondly, there are currently three major *pyguppyclient* releases: i) v0.0.9, which is compatible with guppy v4.4+, ii) v0.0.7a1, which is compatible with guppy v4.0-v4.3 and iii) v0.0.6 (and earlier versions), which are compatible with guppy v3.4-v3.9. These versions are not compatible between them. This implies that if the user wants to use more than one version of *guppy*, the user will need to install multiple versions of *pyguppyclient*. This can be done using Python virtual environments.

Thirdly, the API interface changes slightly between those versions (lack of interface stability). Therefore, that same code that works with one API version may not work with another one. Moreover, we have observed numerous unresolved bugs across various client versions. For example, v0.0.7a1 doesn't return trace and base modification table information (https://github.com/nanoporetech/pyguppyclient/issues/4), while v0.0.9 fails when a modification-aware basecalling model is used (https://github.com/nanoporetech/pyguppyclient/issues/8). While the first issue has been solved in the newer *pyguppyclient* version (v0.0.9), the latter hasn't been addressed so far. Consequently, users cannot retrieve trace (for guppy v4.0-v4.3) and modification tables (for guppy v4.0+) using *pyguppyclient* as of now.

We have addressed all above in *modPhred* by: i) making it compatible with all guppy versions starting from v3.4.1, ii) solving the limitations of the existing client mentioned above (using *guppy_encode_live.py*) , and iii) speeding up *on-the-fly* basecalling ~100x by using batch processing.

**Supplementary Note 2: *Megalodon* data aggregation troubleshooting**

*Megalodon* stores per base modification information in a custom sqlite3 database. While convenient, we have found that this storage system has several limitations. Firstly, the database file is large - typically 50 times larger than the resulting BAM files - which leads to increased storage costs. Secondly, in order to report per-site modification information, the database has to be queried multiple times for each position, each query corresponding to a given read. Because the database is stored in the hard-drive (not in memory), high-speed drives (e.g. SSD) are recommended to perform the database queries. Thirdly, this system seems to be highly inefficient for high coverage samples. For example, we found that the high-coverage (900x) *E. coli* DNA sequencing sample (PRJEB22772) was timed out in the cluster node after 120 hours (**Table S2**). To overcome this limitation, we ran data aggregation locally using SSD drives. However, we found that even this solution was still insufficient, as *megalodon* was unable to finish processing the dataset after 7 days of computing. By contrast, *modPhred* had finished processing the same dataset in less than 6h using the same computing resources (**Table S2**).

### 3. SUPPLEMENTARY TABLES

**Table S1:** Feature comparison: *megalodon* and *modPhred*

| | **megalodon v0.1.0** | **megalodon v2.2.10** | **modPhred v1.0** |
|---|---|---|---|
| **Input** | | | |
| accepts basecalled Fast5 | N | Y [a] | Y |
| accepts raw Fast5 with live basecalling support | Y [b] | Y | Y |
| support for *guppy* < 4.0 | N | N | Y |
| support for *guppy* > 4.0 | N | Y | Y |
| support local basecalling | Y | Y | Y |
| support remote basecalling | N | N | Y |
| DNA support | Y | Y | Y |
| genome alignment for DNA | Y | Y | Y |
| RNA support | N | Y | Y |
| genome (spliced-aware) alignment for RNA | N | N | Y |
| **Running** | | | |
| can process multiple samples at once | N | N | Y |
| can process one sample in batches | N | N | Y |
| modular design | N | N | Y |
| encoding modifications in FastQ | N | N | Y: quals |
| encoding modifications in BAM | N | Y: MM/MP tags | Y: quals |
| **Output** | | | |
| BAM output | N | Y | Y |
| bedMethyl output | Y | Y | Y |
| per-read information | Y: custom db | Y: custom db | Y: quals |
| per-read visualisation | N | N | Y |
| co-occurrence | N | N | Y |
| read clustering | N | N | Y |
| **Dependencies** [c] | | | |
| Guppy Client | Not needed | 0.0.7+ | 0.0.6+ |
| Guppy Basecaller | Not needed | 4.0+ | 3.4+ |
| CUDA | Not needed | 10+ | 10+ |
| taiyaki | Needed | Not needed | Not needed |

[a] Reads need to be basecalled with --post_out parameter.
[b] It uses *taiyaki* backend to basecall reads and retrieve modification information.
[c] See ***Supplementary Note 1*** on Guppy basecaller API troubleshooting

**Table S2.** Performance comparison: *megalodon* and *modPhred*

| | **Megalodon v2.2.10** (guppy v4.0.15) | **modPhred v1.0** (guppy v4.0.15) | **Difference** (fold change) |
|---|---|---|---|
| **Runtime** [a] [hh:mm:ss] | | | |
| test | 0:38:57 | 0:11:42 | 3.33 |
| PRJEB22772 | >124:00:00 [b] | 5:21:05 | > 23.17 |
| PRJNA477598 | 52:51:50 | 13:39:37 | 3.87 |
| **Peak memory usage** [GB] | | | |
| test | 8.27 | 2.37 | 3.48 |
| PRJEB22772 | 16.00 | 9.22 | 1.73 |
| PRJNA477598 | 16.00 | 16.00 | 1.00 |
| **Output size** [GB] | | | |
| test | 3.30 | 0.06 | 55.00 |
| PRJEB22772 | 169.00 | 3.10 | 54.51 |
| PRJNA477598 | 278.00 | 5.70 | 48.77 |

[a] *Runtime computed using GPU NVIDIA GeForce RTX 2080 Ti*
[b] *Megalodon did not finish after 124h, see also **Supp Note 2***

**Table S3**. Metadata of the WGS datasets used for benchmarking *modPhred* and *megalodon*

| | **test set** (subset of PRJEB22772) | **PRJEB22772** | **PRJNA477598** |
|---|---|---|---|
| runs | 2 | 2 | 13 |
| reads | 15,063 | 749,986 | 2,405,041 |
| size [bytes] | 1,992,412 | 103,356,768 | 213,039,176 |
| bases | 136,370,224 | 6,351,379,283 | 11,155,431,907 |
| chemistry | R9.4 | R9.4 | R9.4 |
| device | MinION | MinION | MinION |
| sample type | Genome sequencing | Genome sequencing | Genome sequencing |
| species | *E. coli* (K12) | *E.coli* (K12) | ZymoBIOMICS microbial community standard |

**Table S4.** Predicted DNA-modified sites and their penetrance in microbial datasets using *ModPhred*.

| project | species | depth | modified positions | | | | penetrance | | |
|---|---|---|---|---|---|---|---|---|---|
| | | | **m6A** | **5mC** | **mean m6A freq** | **mean m5C freq** | **m6A GATC** | **5mC CCWGG** | **5mC CpG** |
| PRJNA477598 | *C. neoformans* | 66 | 17,204 | 72,071 | 0.134 | 0.102 | 0.102 | 0.001 | 0.039 |
| | *P. aeruginosa* | 147 | 65,784 | 9,990 | 0.260 | 0.078 | 0.776 | 0.001 | 0.003 |
| | *E. coli* | 347 | 36,912 | 29,488 | 0.634 | 0.370 | 0.939 | 0.938 | 0.004 |
| | *E. coli* | 46 | 39,511 | 43,594 | 0.592 | 0.262 | 0.938 | 0.933 | 0.014 |
| | *E. faecalis* | 375 | 2,052 | 4,441 | 0.125 | 0.081 | 0.135 | 0.000 | 0.019 |
| | *L. monocytogenes* | 196 | 1,982 | 4,669 | 0.126 | 0.079 | 0.156 | 0.001 | 0.017 |
| | *B. subtilis* | 172 | 14,132 | 11,016 | 0.165 | 0.170 | 0.274 | 0.499 | 0.010 |
| | *S. enterica* | 73 | 37,436 | 32,299 | 0.632 | 0.319 | 0.881 | 0.826 | 0.004 |
| | *saureus* | 244 | 79 | 10,269 | 0.089 | 0.150 | 0.003 | 0.000 | 0.046 |
| | *S. cerevisiae* | 87 | 973 | 8,660 | 0.116 | 0.092 | 0.014 | 0.000 | 0.014 |
| PRJEB22772 | *E. coli* | 905 | 38,897 | 29,165 | 0.632 | 0.424 | 0.998 | 0.998 | 0.001 |
| | *E. coli* | 251 | 38,495 | 29,869 | 0.593 | 0.441 | 0.997 | 0.999 | 0.004 |

**Table S5.** Predicted RNA modification levels using *modPhred* in diverse mixes of RNA-modified reads, used to benchmark and illustrate the features of modAnalysis module.

| sample stoichiometry - ground truth [unmod:m6A:m5C:5hmC] | median coverage | Predicted **$m^6A$** modification frequency | Predicted **$m^5C$** modification frequency | Predicted **$hm^5C$** modification frequency |
|---|---|---|---|---|
| 50:30:10:10 | 469 | 30.23 | 10.5 | 9.98 |
| 50:30:15:5 | 469 | 30.24 | 15.48 | 4.99 |
| 50:30:18:2 | 468 | 30.25 | 18.44 | 2.03 |

**Table S6.** Runtime for *modPhred* on test dataset depending on the computing platform and *guppy* version

| | **Guppy version** | |
|---|---|---|
| **Platform** | v3.6.1 | v4.0.15 |
| 6CPUs [a] + local GPU [b]<br>(modPhred with local basecalling) | 0:08:44 | 0:11:42 |
| 6CPUs [c] + remote GPU [b]<br>(modPhred with remote basecalling) | 0:06:33 | 0:05:06 |

[a] *GPU NVIDIA GeForce RTX 2080 Ti*
[b] *CPU remote Intel(R) Xeon(R) Silver 4214 CPU @ 2.20GHz*
[c] *CPU workstation Intel(R) Core(TM) i7-8700 CPU @ 3.20GHz*

**Table S7.** Runtime for *modPhred* on test dataset depending on the number of CPUs

| **Runtime [a]**<br>[hh:mm:ss] | | | |
|---|---|---|---|
| | **Number of CPUs** | | |
| | 6 | 12 | 18 |
| test | 0:11:42 | 0:09:00 | 0:07:17 |
| PRJEB22772 | 5:21:05 | 4:17:42 | 3:42:01 |

[a] Runtime computed using *CPU Intel(R) Xeon(R) Silver 4214 CPU @ 2.20GHz + NVIDIA GeForce RTX 2080 Ti*

**Table S8.** Comparison of *modPhred* runtimes using remote or *on-the-fly* basecalling settings

| Runtime [a] [hh:mm:ss] | | |
|---|---|---|
| | guppy v3.6.1 with --fast5_out | modPhred with *on-the-fly* basecalling |
| PRJEB22772 | 19:13:00 | 5:21:05 |
| PRJNA477598 | 57:29:00 | 13:39:37 |

[a] *Runtime computed using CPU Intel(R) Xeon(R) Silver 4214 CPU @ 2.20GHz + NVIDIA GeForce RTX 2080 Ti*

### 4. SUPPLEMENTARY FIGURES

**Figure S1.** Schematic overview of the functionalities of each of the 4 modules included in *modPhred*: (i) *modEncode*, (ii) *modAlign*, (iii) *modReport* and (iv) *modAnalysis*.

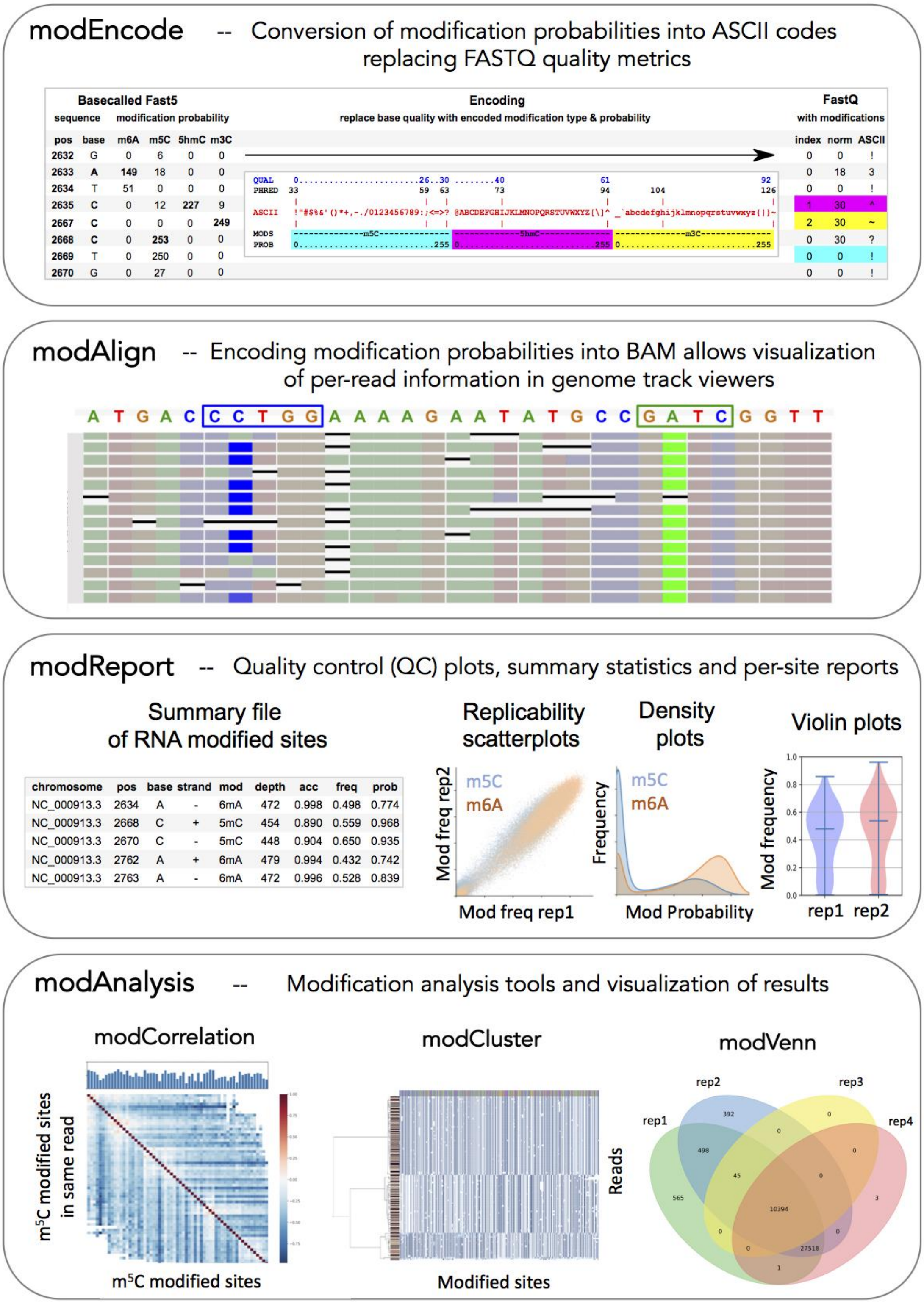

**Figure S2. Analysis of DNA and RNA modifications using modPhred. (A)** Reproducibility of *E. coli* DNA modifications predicted using *modPhred*. Only sites modification frequency greater than 0.05 are considered as 'modified' and included in the analysis. **(B)** Density plots depicting the reproducibility of modification frequencies at individual sites across biological replicates (upper panels). We should note that the reproducibility is very high despite the sequencing depth being relatively distinct across the two biological replicates (bottom panel). **(C)** General quality control statistics generated by modReport module from modPhred. modReport supports both DNA and RNA modifications. **(D)** Hierarchical clustering of reads based on their RNA modification patterns, using modClustering script which is part of the modAnalysis module in ModPhred. *ModPhred* also computes the number of optimal read clusters using Scree's plot.

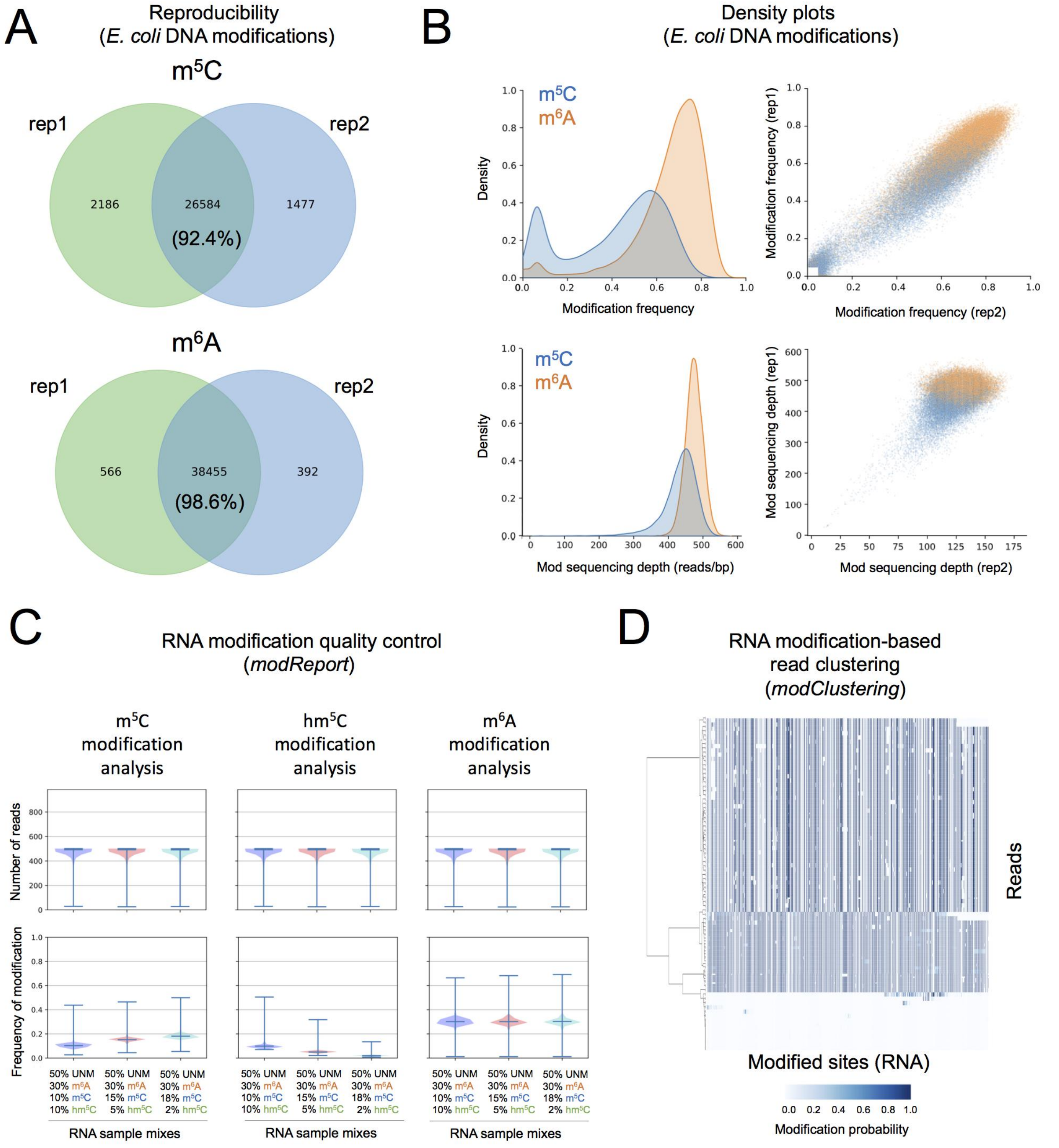

**Figure S3**. **Comparison of ModPhred and megalodon in DNA modification detection. (A)** Comparison of $m^6A$ and $m^5C$ DNA modification predictions obtained using *megalodon* (blue) and *modPhred* (green) in the *E. coli* whole genome sequencing dataset (PRJEB22772). *ModPhred* will only consider a site as 'modified' if the modification frequency at a given site is greater than 0.05. **(B)** Comparison of the $m^6A$ and $m^5C$ DNA modification frequencies (proportion of modified reads relative to stranded coverage) between megalodon and *ModPhred*.

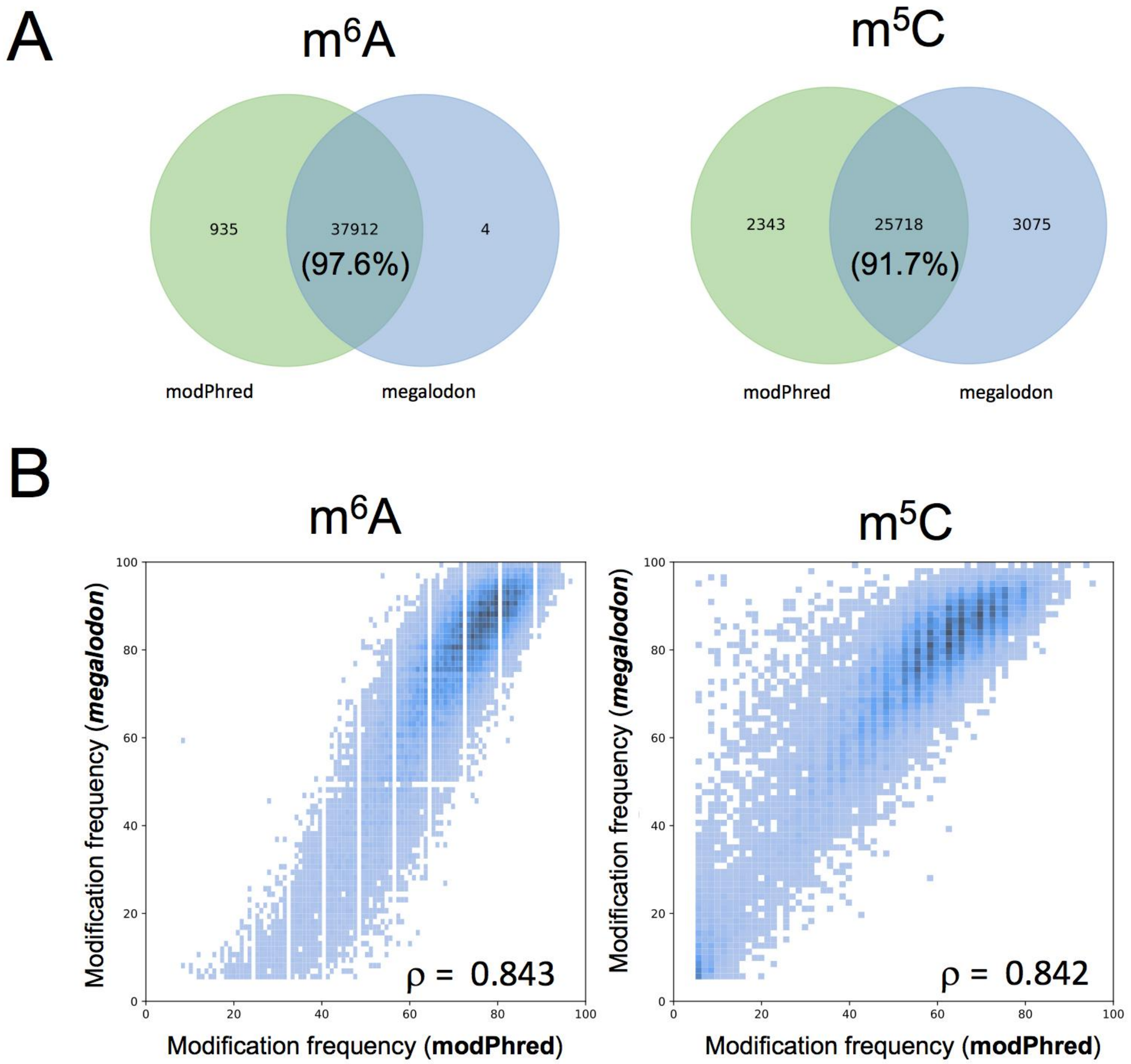
